# Molecular recording of mammalian embryogenesis

**DOI:** 10.1101/384925

**Authors:** Michelle M. Chan, Zachary D. Smith, Stefanie Grosswendt, Helene Kretzmer, Thomas Norman, Britt Adamson, Marco Jost, Jeffrey J. Quinn, Dian Yang, Alexander Meissner, Jonathan S. Weissman

**Affiliations:** Department of Cellular and Molecular Pharmacology, University of California, San Francisco, San Francisco, CA, USA; Howard Hughes Medical Institute, University of California, San Francisco, San Francisco, CA, USA; Center for RNA Systems Biology, University of California, San Francisco, San Francisco, CA, USA; Department of Microbiology and Immunology, University of California, San Francisco, San Francisco, CA, USA; Broad Institute of MIT and Harvard, Cambridge, Massachusetts, USA; Department of Stem Cell and Regenerative Biology, Harvard University, Cambridge, Massachusetts, USA; Department of Molecular and Cellular Biology, Harvard University, Cambridge, Massachusetts, USA; Department of Genome Regulation, Max Planck Institute for Molecular Genetics, Berlin 14195, Germany

## Abstract

Understanding the emergence of complex multicellular organisms from single totipotent cells, or ontogenesis, represents a foundational question in biology. The study of mammalian development is particularly challenging due to the difficulty of monitoring embryos *in utero*, the variability of progenitor field sizes, and the indeterminate relationship between the generation of uncommitted progenitors and their progression to subsequent stages. Here, we present a flexible, high information, multi-channel molecular recorder with a single cell (sc) readout and apply it as an evolving lineage tracer to define a mouse cell fate map from fertilization through gastrulation. By combining lineage information with scRNA-seq profiles, we recapitulate canonical developmental relationships between different tissue types and reveal an unexpected transcriptional convergence of endodermal cells from extra-embryonic and embryonic origins, illustrating how lineage information complements scRNA-seq to define cell types. Finally, we apply our cell fate map to estimate the number of embryonic progenitor cells and the degree of asymmetric partitioning within the pluripotent epiblast during specification. Our approach enables massively parallel, high-resolution recording of lineage and other information in mammalian systems to facilitate a quantitative framework for describing developmental processes.

## Introduction

Development of a complex multicellular animal from a single cell is an astonishing process. Classic lineage tracing experiments using *C. elegans* as a model revealed surprising outcomes, including deviations between lineage and functional phenotype, but nonetheless benefited from the highly deterministic nature of this organism’s development^1^. Alternatively, higher-order organisms generate larger, more elaborate structures that progress through multiple, stochastically regulated transitions, raising questions regarding the timing and coordination between specification and commitment to ensure faithful recapitulation of an exact body plan^2,3^.

Single cell RNA-sequencing (scRNA-seq) has permitted unprecedented explorations into cell type heterogeneity and correspondingly renewed interest in cell fate mapping in the form of trajectory inference and cell state landscape reconstruction^4^. Studies profiling tens of thousands of cells have uncovered transcriptional programs and inferred cellular relationships underlying numerous biological processes including the development of flatworms^5,6^, frogs^7^, zebrafish^8,9^, and mouse^10,11^. More recently, CRISPR-Cas9-based technologies have made direct recording of cell lineage possible^12–14^ and, in combination with scRNA-seq, have been applied in zebrafish to elucidate the fate map underpinning embryogenesis, brain development, haematopoiesis, and fin regeneration^15–17^. However, previous implementations of lineage tracing have relied on one or two bursts of barcode diversity generation, which is well suited for those studies but may limit them in more complex applications, such as monitoring mammalian development. Indeed, to date the feasibility of these approaches for the generation of a mammalian fate map remains unresolved.

An ideal molecular recorder for these questions would possess the following characteristics: 1) minimal impact on a cell’s behaviour or developmental trajectory; 2) high information content to report the relationships between hundreds of thousands of cells precisely; 3) a single cell read out for simultaneous profiling of functional state^15–17^; 4) flexible recording rates that can be tuned to the developmental process in question, ranging from rapid transitions over the course of days to slower processes that unfold over months; and 5) continuous generation of diversity throughout the duration of the monitored process. The last point is especially relevant for recording in mammalian development, where spatial plans are more gradually and continuously specified and may originate from small, transient progenitor fields that could create bottlenecks of variable sizes. Moreover, scRNA-seq in mammalian systems have revealed populations of cells with a continuous spectrum of phenotypes, implying that differentiation does not occur instantaneously or as discrete step-wise transitions and further motivating the need for an evolving molecular recorder^4^.

Here, we generated and validated a method for simultaneously reporting cellular state and lineage history for thousands of individual cells from single mouse embryos. Our CRISPR-Cas9-based molecular recorder is capable of high information content and multi-channel recording with readily tunable mutation rates. We employ the recorder as a continuously evolving lineage tracer to observe the fate map underlying mouse embryogenesis through gastrulation and after the body pattern has been set, recapitulating canonical paradigms for cell type commitment and illustrating how lineage information may facilitate the identification of novel cell types.

## Results

### A transcribed, multi-channel, and continuously evolving molecular recorder

We set out to create a DNA-based molecular recorder capable of capturing high information content from multiple independent signals, while being sufficiently flexible to accommodate a wide range of time scales. Similar to recent reports^12–17^, we utilized Cas9 to generate insertions or deletions (indels) upon repair of double-stranded breaks, which are inherited in the next generation of cells. We record within a 205 base pair, synthetic DNA “target site” containing three cut sites, as well as an 8 base pair randomer to serve as a static “integration barcode” (intBC), which we deliver into cells in multiple copies via piggyBac transposition (**Fig. 1a, b**). The target site is embedded in the 3’UTR of a fluorescent protein under control of a constitutive Ef1α promoter to enable profiling from the transcriptome. A second vector encodes three guide RNAs, each under the control of an independent promoter, to permit recording of multiple, distinct signals (**Fig. 1a, b**)^18^.

**Figure 1:**
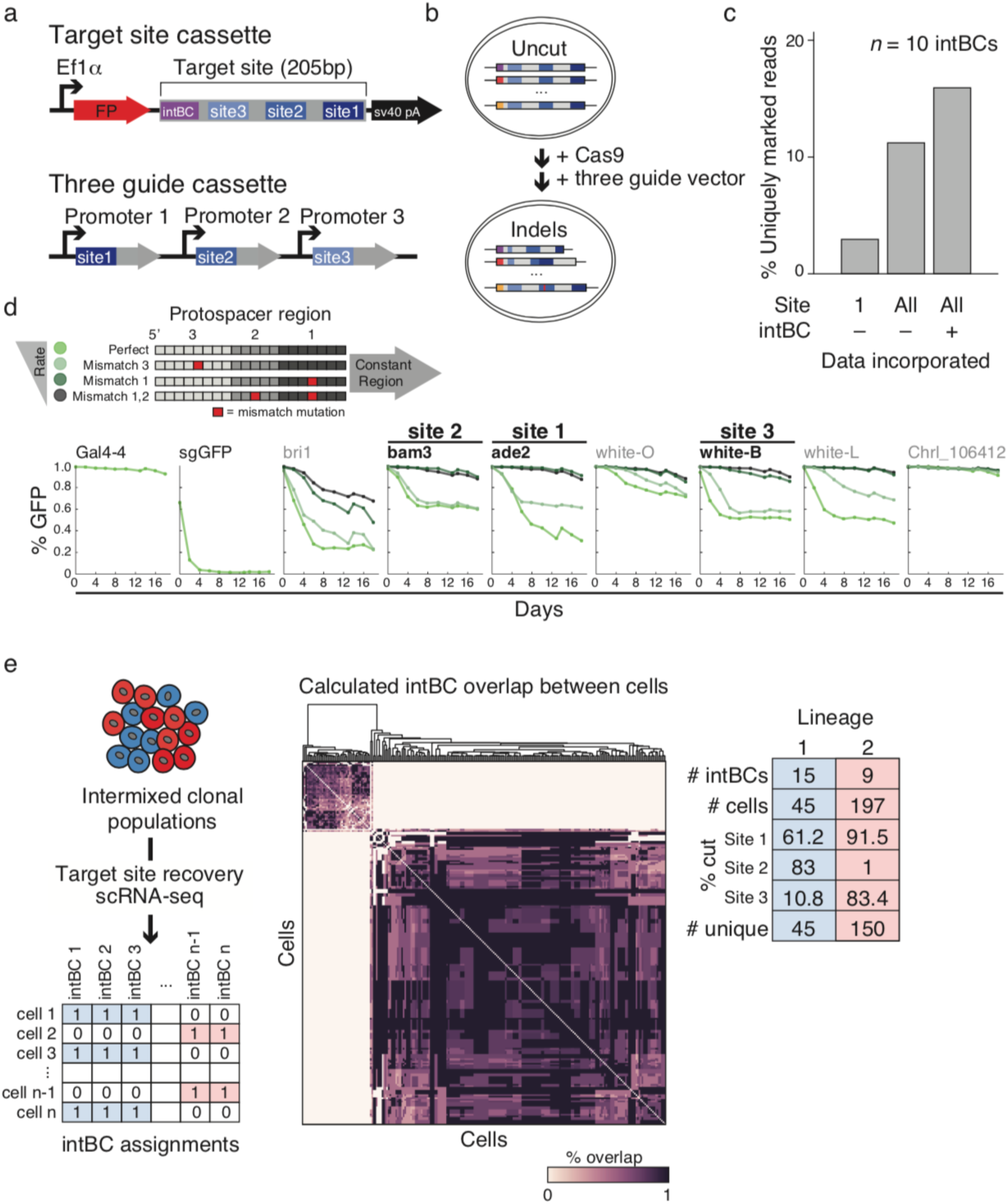
Optimization of a multi-purpose molecular recorder. a. Schematic of the target site and three guide cassettes used in this study. The target site consists of an integration barcode (intBC) and three protospacer sequences for mutation-based recording by Cas9. The target site is located in the 3’UTR of a fluorescent protein (FP) under the control of a constitutive promoter, Ef1. The three guide cassette encodes three different single guide RNAs (sgRNAs), each under the control of an independent promoter. A combination of constitutive and responsive promoters can be used to permit recording of multiple signals. For applications described here, three constitutive promoters (mU6, hU6, and bU6) are used. b. Schematic of molecular recording. Each cell contains multiple genomic integrations of the target site, which are distinguished by their intBCs (left most colored box in target sites). sgRNAs expressed from the three guide cassette direct Cas9 to their cognate protospacer sequences on the target to generate insertion (red) or deletion mutations. c. Information content scales according to the number of sites used and the presence of the intBC. In this experiment, a population of K562 cells with 10 target site integrations was subjected to molecular recording by Cas9 for 6 days and the percentage of uniquely marked reads was calculated using the following information: site 1 only with intBCs masked, sites 1-3 (All) with intBCs masked, and sites 1-3 (All) with intBCs considered. By incorporating multiple protospacers within identifiable cassettes, more reads can be distinguished from each other to maximize data recovery from the system. d. Guide RNA mismatches to the target site alter mutation rate. A DNA sequence containing seven independent protospacers was incorporated into the 5’ coding sequence of a single copy GFP reporter to measure the mutation rate. The protospacer-recognizing sequence of an sgRNA can be separated into three adjacent regions by proximity to the PAM: region 1 (proximal), region 2, and region 3 (distal). sgRNAs with either single mismatches to these regions or dual mismatches in 1 and 2 were integrated into a population of reporter-expressing cells and the fraction of GFP+ cells was measured by flow cytometry over a 20 day time course. An sgRNA against Gal4-4 and the coding sequence of GFP were used as negative and positive controls, respectively. Cut sites are ordered as they appear in the reporter from 5’ to 3’. The ade2, bam3, and white-B protospacers were selected for incorporation into the final target site to take advantage of the dynamic range of cutting efficiencies generated by their respective sgRNA series. e. Simultaneous and continuous molecular recording of multiple clonal populations in K562 cells. (Left) Strategy for partitioning a multi-clonal population into their clonal populations. Target sites are sequenced from a scRNA-seq library (See **Fig. S2**) and the intBCs in each cell is identified as present or absent. (Middle) Heatmap of the percent overlap of intBCs between all cells. The cells segregate into two populations representing the descendants of two progenitor cells from the beginning of the experiment. (Right) Table summarizing results of the experiment, including the generation of indels over the experiment duration.

Our system is capable of high information storage due to the large diversity of heritable Cas9 repair outcomes, the utilization of many cut sites, and the presence of an intBC to distinguish between different genomic integrations of the target site. While DNA repair is able to generate hundreds of unique indels, the indel distribution is different for each guide and not uniform: some guides produce highly biased outcomes while others create a diverse indel series (**Fig. 1c**, **Fig. S1**)^19–21^. Indeed, the number of uniquely marked reads in a population of mutated target sites is lower when the integration barcode is not taken into account (**Fig. 1c, Fig. S1**).

The ability to tune the mutation rate of the molecular recorder is critical for profiling processes that span time scales that are not pre-defined, but range from days to weeks to months. To identify sequences with a broad range of cutting rates, we screened several protospacers by modifying the complementarity of each guide^22^, and observed the delayed disruption of a GFP reporter over a 20-day time series (**Fig. 1d**). Slower cutting rates may also improve viability *in vivo*, as frequent double-strand breaks resulting from Cas9 activity may cause cellular toxicity leading to a fitness cost.

Our integration barcoding strategy also permits tracking multiple discrete founder populations simultaneously, which we tested by stably transducing K562 cells with a high complexity library containing thousands of unique intBCs. Because the likelihood of two cells integrating the same set of intBCs is small, the complement of intBC sequences distinguishes individual progenitors. To illustrate this point, we sorted populations of 10 cells and propagated them for 18 days. We utilized the 10x Genomics platform to generate a primary, cell-barcoded cDNA pool that can be split into two fractions: one to generate a global transcription profile and the other to specifically amplify our target site (**Fig. S2**). We detected two populations of cells by their integration barcodes, implying that only two clonal lineages expanded from the initial population of 10 (**Fig. 1e**). Moreover, each population exhibited mutations in their target sites, showcasing our ability to combine dynamic recording with traditional static lineage tracing.

### Tracing cell lineages in mouse development

With a method for molecular recording established, we applied our technology to generate a cell fate map of mouse early development from totipotency onwards. We integrated multiple target sites into the zygotic genome and initiated cutting by the delivery of a paternally expressed Cas9-GFP transgene, which is constitutively expressed following zygotic genome activation (**Methods**). We screened blastocyst-stage embryos for expression of the target site using mCherry, transferred highly fluorescent embryos into pseudopregnant females, and isolated them after six or seven days of internal gestation at ~embryonic day (E)8.5 or E9.5 for lineage analysis (**Fig. 2a**). To confirm our molecular recorder’s ability to act as a lineage tracer, we amplified the target site from bulk tissues representing the placenta, yolk sac, and three embryonic fractions from a single E9.5 embryo (**Fig. 2b**). We were able to recapitulate expected tissue relationships by clustering tissues according to the similarity in their indel proportions, with the placenta forming the outgroup, followed by yolk sac, and with the three embryonic tissues showing the most similar lineage relationships (**Fig. 2b**).

**Figure 2:**
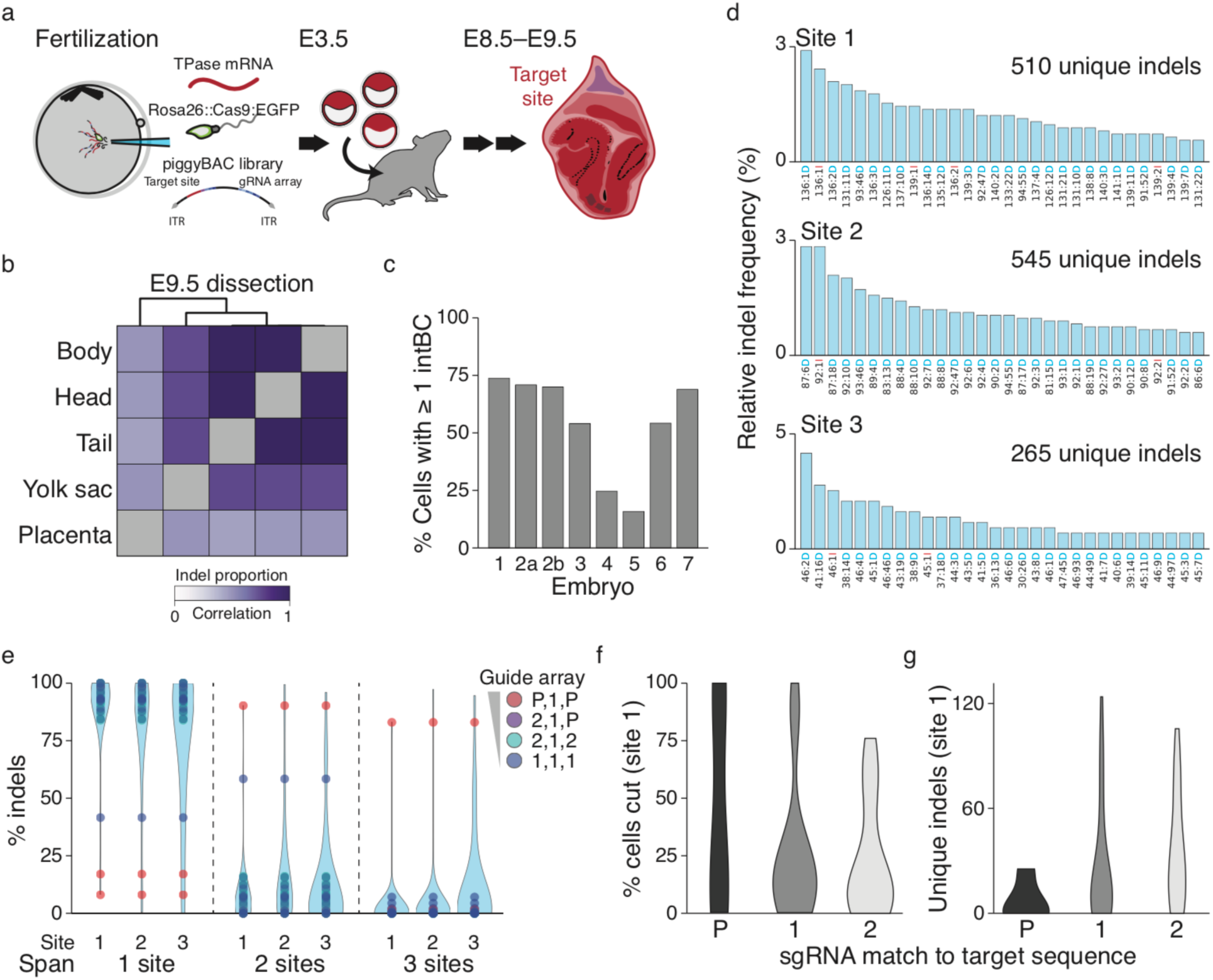
Lineage tracing in mouse from fertilization through gastrulation. a. Schematic of lineage tracing in mouse experiments. The target site (in the 3’UTR of mCherry) and the three guide cassettes are encoded in one vector flanked by piggyBac transposon inverted terminal repeats (ITRs) to permit multiple genomic integrations. The vector, along with piggyBac transposase mRNA, and Rosa26::Cas9:EGFP strain sperm, are injected into oocytes to enable lineage tracing in all cells. Embryos are cultured *in vitro* until the blastocyst stage (E3.5), screened for high expression of the target site vector, and transferred into pseudo-pregnant females. Embryos were isolated at E8.5 for single cell analysis or E9.5 for bulk tissue analysis. b. Heatmap of correlation coefficients between indel proportions recovered from dissected, bulk tissue of an E9.5 embryo. Placenta is the most distantly related, followed by the yolk sac, with the three embryonic compartments sharing the highest similarity, recapitulating their known developmental relationships. c. The percentage of cells with at least one target site sequenced from seven embryos. The target site is separately recovered from the primary cell-barcoded cDNA pool, while the total number and transcriptional identity of cells is generated from the scRNA-seq library (see **Fig. S2**). d. Distribution of relative indel frequencies estimated from 40 independent target sites taken from all embryos. Each site produces hundreds of outcomes, showing the potential for high information encoding. The indel code along the x-axis is as follows: “Alignment Coordinate: Indel Size Indel type (**I**nsertion or **D**eletion).” e. Violin plots representing the distribution of indels that span one, two, or three sites, shown per site. Indels that span more than one site indicate large deletions, which can disrupt additional recording. However, most instances of large deletions appear to result from simultaneous double strand breaks, as the number of multi-site-spanning indels are substantially reduced by using slower cutters. Colors indicate the three guide array used, with the mutation to the target sequence for each included (see **Figure 1d**). P = no mismatches, 1 = mismatch in region 1, 2 = mismatch in region 2 f. The percentage of cells carrying mutations decreases according to the mismatch between the protospacer and guide. All cells in an embryo are related to each other. Thus, mutations which happen earlier during development are propagated to more cells, leading to a higher percentage of cells with indels. g. Indel diversity is inversely related to cutting efficiency. For site 1, the number of unique indels were counted according to the protospacer-guide mismatch. Mutations that occur earlier in development are propagated to more cells leading to a smaller number of unique indels.

With this *in vivo* proof of principle established, we proceeded to generate single cell data from additional embryos (**Fig. S2**). We selected E8.5 as our earliest recovery stage because it allows for relatively deep coverage of single embryos, but also follows the determination of major extra-embryonic and embryonic compartments, including many primary derivatives of the embryonic germ layers. Total cell counts for seven embryos, including yolk sac and placenta, ranged between 13,680 – 56,400 cells, with 7,364 – 12,990 cells (15.8% – 61.4%) profiled by scRNA-seq and encoding 167 – 2,461 unique lineage identities. After dissociation of the embryo, we sampled embryo 2 twice, with each sample occupying independent 10x lanes to act as technical replicates. We capture at least one target site in 15% – 75% of cells and recover between 3 and 15 intBCs, though many target sites are either lowly or heterogeneously represented (**Fig. 2c, Fig. S3**). We reasoned that incomplete target site capture in embryos may be due to lower expression and changed the promoter from a truncated form of Ef1α which functions well in the human K562 cell line, to an intron-containing version that improves transcript abundance^23^. After repeating the injection, the resulting embryo (embryo 7) indeed exhibited ~5x higher target site UMI counts and increased capture frequencies (**Fig. S3**). Using 40 independent target sites taken from all seven embryos, we find that many indels are shared with K562 cells, though their likelihoods are different, suggesting that cell type or developmental status may influence repair outcomes (**Fig. 2d**, **Fig. S1, S4**)^19^. Our ability to independently measure and control the rate of cutting across the target site is largely preserved *in vivo*, with minimal interference between cut sites except when using combinations of the fastest cutting guides that likely reflects end-joining between simultaneous double strand breaks (**Fig. 2e**). As expected, the fastest cutters result in a higher proportion of cells with indels compared to slower cutters, indicative of mutations made earlier in development, and this comes with a corresponding reduction in indel diversity (**Fig. 2f, g**). Importantly, at E8.5, most embryos still have unmodified cut sites, which shows that the lineage tracer retains additional recording capacity beyond the temporal interval studied here (**Fig. 2f**).

### Assigning cellular phenotype by simultaneous scRNA-seq

Next, to assign functional identities to cells from our lineage-traced embryos, we utilized annotations from an independent compendium of wild-type mouse gastrulation (E6.5 – E8.5), which identified lineage markers for many tissues, including placental (Elf5, Gata2, Tfap2c), visceral and definitive endoderm (Hnf4a, Apoa4, Afp or Sox17 and Foxa2, respectively), primordial germ cells (Dppa3, Pou5f1/Oct4), neural ectoderm (Otx2, Sox1), surface and placodal ectoderm (Dlx5, Krt8), amnion (Foxf1), blood (Klf1, Gata1), and heart (Nkx2-5). In addition to tissue specificity, the annotations carry information about developmental timing, for example by distinguishing between early and late visceral endoderm compartments that are predominant in E8.0 vs E8.5. We assigned cells from our lineage-traced embryos according to their proximity to the mean expression signature of each cluster and aged each embryo according to their tissue proportions compared to each stage (**Fig. 3a-d**). Notably, we identify primordial germ cells (PGCs) in all embryos, despite their rarity during this moment in developmental time (<0.2% of cells sequenced). Six of our embryos appear to be morphologically normal and contain cells of every expected tissue type. Two of the six embryos mapped to E8.5, and the remaining four embryos mapped to E8.0, though these are more likely to be between E8.0 and E8.5 as the proportions of caudal lateral epiblast and allantois are smaller than expected for E8.0 and indicative of later developmental timing (**Fig. 3d, Fig. S5**). Placenta was not specifically included during embryo isolation and is present in four of six embryos, but serves as a valuable outgroup in these samples to establish our ability to track developmental transitions to the absolute earliest bifurcation. We conclude from these observations that potential fitness costs of double strand break generation and random integration of the transposons in our system do not substantially perturb these early developmental processes beyond what may be a subtle delay in embryo progression.

**Figure 3:**
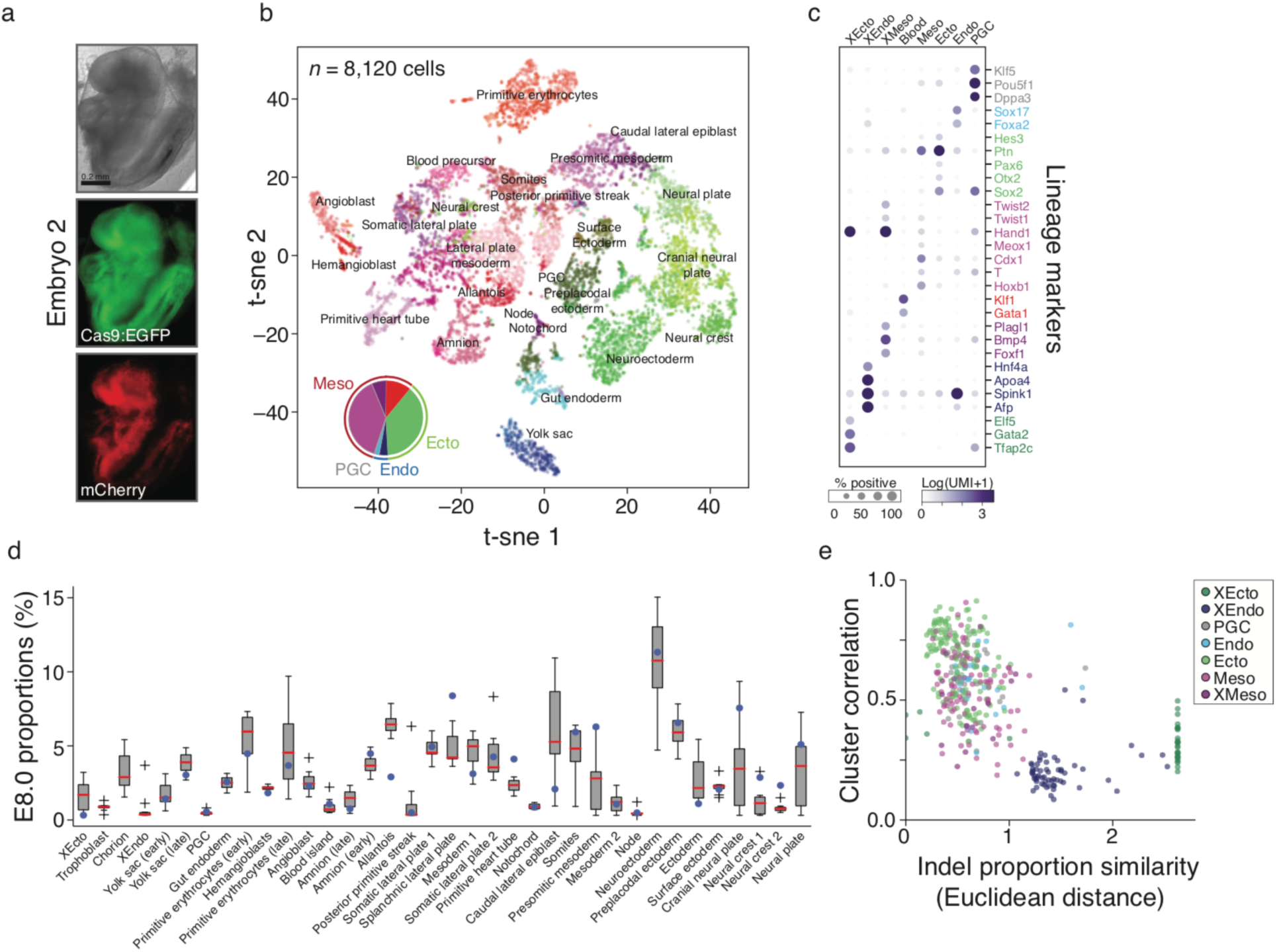
Assigning cellular phenotype by scRNA-seq. a. Image of a lineage-traced E8.5 embryo (embryo 2, see **Fig. S2**) generated using a three guide array in which each carries a protospacer mismatch in region 1. Expression of Cas9:EGFP and the mCherry target site reporter is shown. b. Plot (t-sne) of scRNA-seq from embryo in **a** with corresponding tissue annotations. (Inset) Pie chart of the proportions for different germ layers. For each germ layer, lighter shades refer to embryonic components while darker shades refer to extra-embryonic components. Mesoderm is further separated to include blood (red). c. Dot plot summarizing the expression pattern for canonical tissue-specific markers used in cluster assignment. The fraction of marker-positive cells within the cluster is indicated by the size of the circle, while the color intensity refers to the normalized expression level (cluster mean). At this level of cell-state clustering, the fraction of cells from germ layers with diverse tissue types, such as ectoderm and mesoderm, are smaller, but the specificity of those markers to their respective states remains high, especially when considered in combination. XEcto, extra-embryonic ectoderm/placenta; XEndo, extra-embryonic endoderm/yolk sac; PGC, primordial germ cell; Endo, embryonic endoderm; Ecto, embryonic ectoderm; Meso, embryonic mesoderm; XMeso, extra-embryonic mesoderm. d. Relative tissue proportions from a lineage-traced embryo fall within wild type boundaries. Boxplot for tissue proportions from 10 wild type E8.0 embryos, with the tissue proportions for the embryo in **a** overlaid as blue dots. We compared the tissue proportions for each lineage-traced embryo to each stage of gastrulation from a wild-type compendium with 0.5 day resolution. Mapping to E8.0 represents a slight developmental delay for lineage-traced embryos, which are recovered at ~E8.5. e. Scatter plot showing the relationship between lineage and expression. Cells are assigned to eight early developmental lineages and compared according to their transcriptional similarity and lineage distance (calculated as the Euclidean distance between indel proportions). Colors are assigned in a nested fashion, with all comparisons against XEcto, colored dark green, all comparisons against XEndo, except for XEcto, colored dark blue, etc.

To relate different cell types according to their developmental history, we grouped our clusters according to their germ layer or extra-embryonic vs. embryonic status and characterized their lineage similarity. As the largest and most complex germ layer, we partitioned embryonic mesoderm into its extra-embryonic portion (derived from the posterior primitive streak to form the amnion and allantois) and blood for greater resolution. To infer shared history, we characterized each group by its indel composition (the proportion of each indel) and compared these by Euclidean distance (**Fig. 3e**). When present, placenta (“XEcto”) is the most distantly related tissue, consistent with trophectodermal commitment during early preimplantation development, followed by the visceral endoderm (XEndo), which derives from the second differentiation event within the maturing inner cell mass. PGCs intermix with the embryonic clusters, but often appear more distantly as an embryonic outgroup, consistent with their positional specification, sequestration during early gastrulation, and small founding population. Given that our lineage tracer is able to recapitulate known tissue relationships, we proceeded to investigate specific lineages at single cell resolution.

### Single cell lineage reconstruction of mouse embryogenesis

To analyze our data at single cell resolution, we developed a phylogenetic reconstruction strategy suited to the characteristics of the lineage tracer, namely the categorical nature of indels, the irreversibility of mutations, and the presence of missing values. Briefly, the root node of the tree is established as a vector of length *n* with no mutations, where *n* is the number of cut sites in the embryo (i.e., three times the number of independent target sites, distinguished by their intBCs). An indel from the data set is chosen according to its normalized likelihood (fraction of cells with the indel divided by its estimated indel frequency, see **Fig. 2d**) to establish a new node from which all cells that carry that mutation descend. The process is repeated until no cells compatible with the lineage remain, after which the algorithm traverses up the branch to find more mutations and cells until all are assigned to nodes in the tree. Cells with missing values are assigned to the node that best describes them. To score each tree, we estimated the total tree likelihood by summing the log-likelihoods for all indels that appear in the tree. We simulated at least 40,000 trees for each embryo and chose the highest likelihood tree for downstream analysis (**Fig. 4a,b**, selected from over 100,000 trees). To establish confidence in tree reconstruction, we investigated the consistency of cells below each node across similarly high scoring trees. We find that nodes that represent more cells (ie. correspond to mutations which occur earlier in development) are likely to be consistent, with 31% of all nodes showing perfect consistency (**Fig. S6**). Moreover, functional restriction is apparent when cell type identity from scRNA-seq data is overlaid onto the tree, with fewer cell types as we move from the root to the leaves (**Fig. 4c**).

**Figure 4:**
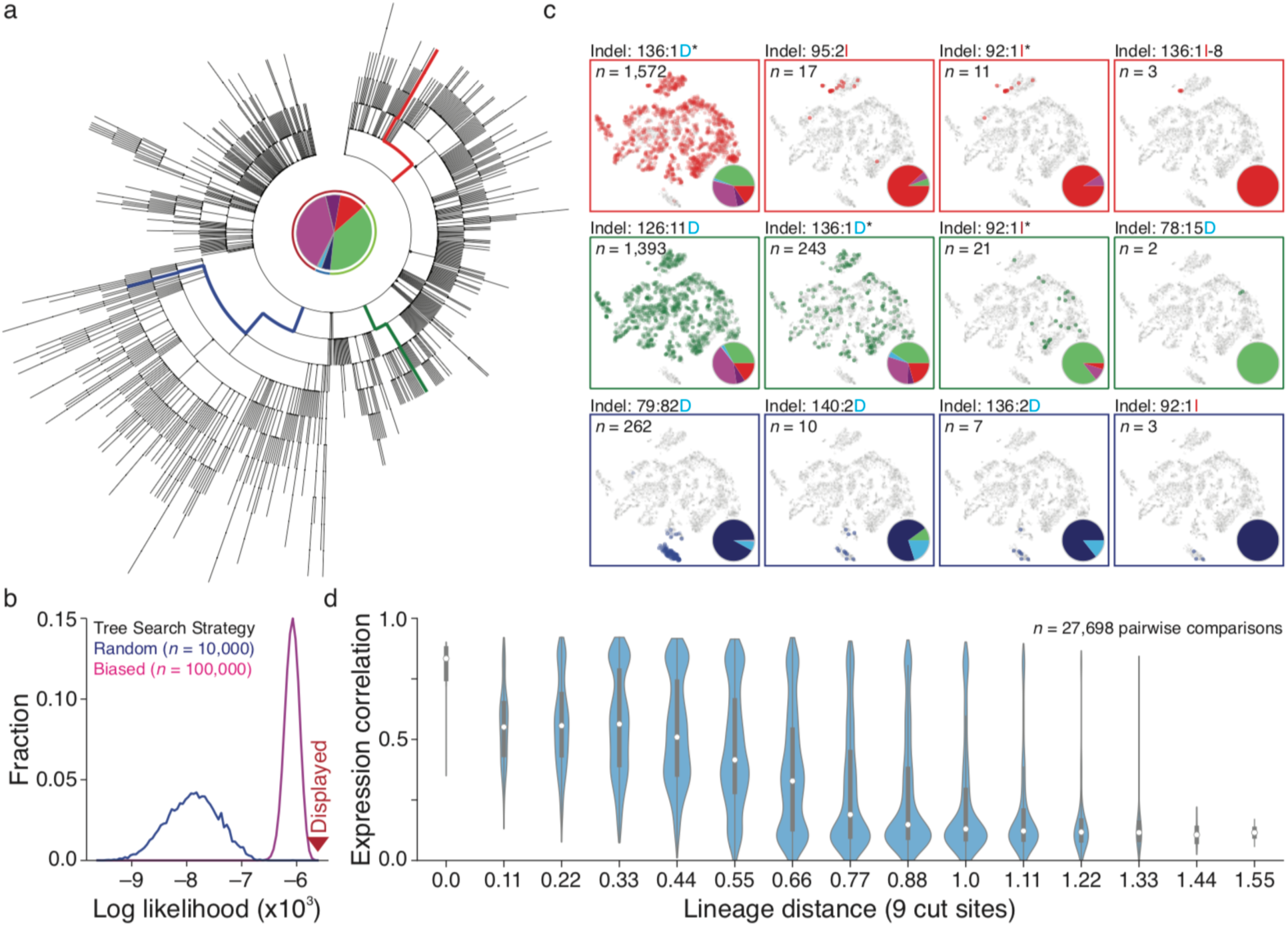
Single cell lineage reconstruction of mouse embryogenesis. a. Reconstructed lineage tree for embryo 2 with three different developmental lineages highlighted. Central pie chart depicts germ layer proportions as in **Figure 3b**. b. Log-likelihood distribution for reconstructed trees. Tree likelihoods are calculated using relative indel frequency estimates (See **Figure 2d**) and the highest likelihood tree is selected for further analysis. Two tree reconstruction strategies are shown. In the random strategy, mutations are sampled randomly from the data set while in the frequency normalized weighted (FNW) strategy, mutations are chosen to bias the search towards higher likelihood trees (See **Methods**). c. Example paths from root to leaf from the selected tree (highlighted by color). Cells for each node in the path are overlaid onto the t-sne representation from **Figure 3b**, with the tissue proportion at each node in the tree included as a pie chart. The number of tissues decreases with higher tree depth, confirming that functional restriction progresses over the course of development. Asterisk, instance of indel predicted to occur more than once for the same intBC from our tree reconstruction. Note that these indels are among the most frequent indels for their sites (see **Figure 2d**). d. Violin plots representing the relationship between lineage and expression for individual pairs of cells. Lineage distance is calculated using a modified Hamming distance. Expression correlation decreases with increasing lineage distance, showing that closely related cells are more likely to share function.

scRNA-seq-based strategies for ordering cells, such as trajectory inference, typically assume that functional similarity is a result of close lineage^4^. To investigate this question directly, we compared cell-to-cell lineage distance to cell-to-cell RNA-seq correlation, using a modified Hamming distance normalized to the number of shared cut sites to measure lineage distance. Generally, cells separated by a smaller lineage distance have more similar transcriptional profiles, though the relationship is clearer in some embryos than in others (**Fig. 4d, Fig. S7**). This result is consistent with the notion of continuous restriction of potency as cells differentiate into progressively restricted cell types.

### Transcriptional state and developmental origin do not always correspond

To systematically explore the lineage relationships across tissues, we grouped cells according to their function as assigned by scRNA-seq, counted the number of shared progenitors between each tissue type in our tree, and clustered the resulting matrix. The reconstructed tissue relationships generally recapitulated canonical knowledge of development and are consistent across independent embryos (**Fig. 5a, Fig. S7**). Nonetheless, there were two tissue pairs that were unexpectedly closely related: the extra-embryonic and embryonic endoderm and the mesoderm and ectoderm. Examination of more refined cluster identities indicates that the close link between mesoderm and ectoderm may be driven by a shared heritage between presomitic mesoderm and neural tissues, corroborating a shared origin for some cells in these clusters from a neuromesodermal progenitor^24^.

**Figure 5:**
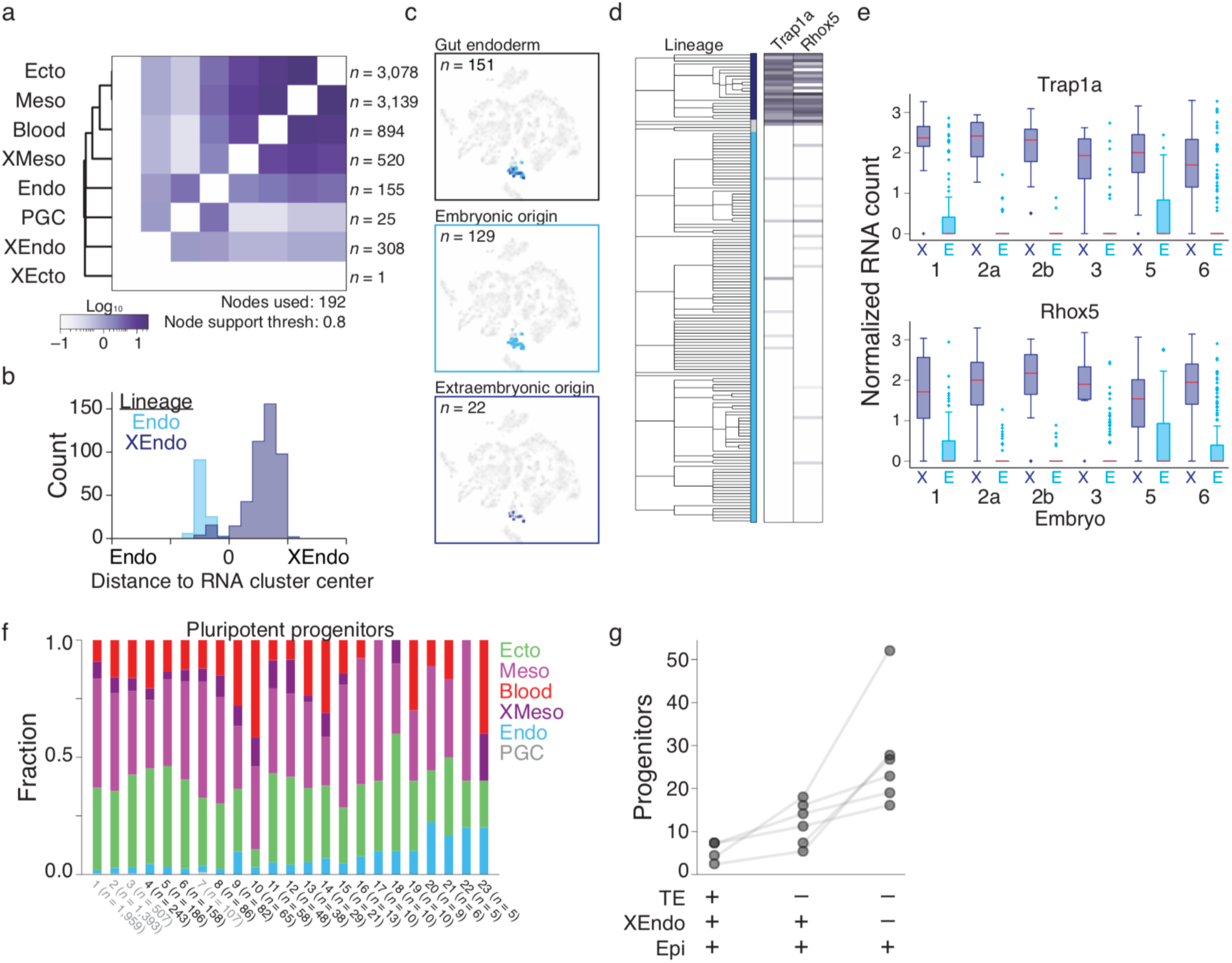
Disparities between transcriptional identity and lineage history within the extra-embryonic endoderm. a. Heatmap of the shared progenitor score for different tissues from embryo 2, calculated as the number of nodes that are shared between two tissues scaled by the number of tissues represented in each node (see **Methods**). Reconstructed tissue relationships using these progenitor scores correctly recreate expected relationships from studies of development. The node support threshold represents the minimum frequency for which an included node must be observed across similarly high scoring trees (see **Fig. S6)**. Heatmaps for additional embryos are included in **Fig. S7**. b. Histograms representing the relative distances from the mean expression profile of either the endoderm cluster or the extra-embryonic endoderm cluster for cells of either embryonic origin (Endo) or extra-embryonic origin (XEndo) for embryo 2. While the majority of XEndo cells cluster together, there is a subset of cells with gut endoderm functional identity that stem from branches with clear extra-embryonic origin. c. Plots (t-sne) of scRNA-seq data for embryo 2, with gut endoderm cells highlighted. Endoderm cells segregate from the rest of the embryo, and cannot be distinguished by origin. d. Lineage tree for endoderm cells from embryo 2 showing the differential expression of two extra-embryonic marker genes, which correspond to cellular origin (middle bar). Each gene is significantly different between the lineages using the Kolmogorov-Smirnov test. Dark blue, extra-embryonic origin; light blue, embryonic origin; grey, ambiguous origin. e. Boxplots for genes Trap1a and Rhox5 confirms consistent differential expression across lineage-traced embryos according to their embryonic or extra-embryonic ancestry. f. Relative tissue distribution of cells descended from reconstructed or profiled pluripotent progenitor cells for embryo 2. “Profiled” refers to a unique lineage identity corresponding to multiple cells directly observed in the data set. While pluripotent cells are able to form all germ layers, they show divergent propensities towards different cell fates, illustrating the asymmetric partitioning that can occur during positional specification from a field of otherwise equivalently potent cells. Pluripotent nodes highlighted in grey represent those that give rise to primordial germ cells. Color assignments are as in **Figures 3** and **4**. g. Estimated progenitor field sizes for three types of early developmental potency. Totipotent cells can give rise to all cells in the developing embryo, including placental lineages of trophectodermal (TE) origin, while pluripotent progenitors are partitioned into early and late modes by their ability to generate extra-embryonic endoderm (XEndo) in addition to the epiblast (Epi).

We next explored the unexpected relationship between extra-embryonic and embryonic endoderm in greater detail. The extra-embryonic endoderm derives from the hypoblast in the second differentiation event in mammalian development, while the embryonic endoderm emerges several days later from the embryo-restricted epiblast. Manual inspection of the trees revealed a small proportion of cells that appear transcriptionally as embryonic endoderm but that lineage analysis shows as present within extra-embryonic branches (**Fig 4c, blue**). Several hypotheses may account for this outcome: 1) cells were assigned to an incorrect type; 2) the lineage reconstruction is incorrect; or 3) cells from multiple origins can have convergent endoderm functional signatures. To investigate the first hypothesis, we compared the expression profiles of these cells to the RNA-seq cluster centers of the extra-embryonic and embryonic endodermal types. The endoderm-classified cells deriving from extra-embryonic origin are correctly assigned to endoderm cell type, but do share slightly higher expression similarity with yolk sac than other endoderm cells (**Fig. 5b, Fig. S8**). Notably, these differences are not obvious when the embryonic or extra-embryonic lineages are overlaid onto the t-sne projection of the full embryo (**Fig. 5c**).

If the lineage reconstruction is correct, i.e. hypothesis 2 is not supported, then we might expect a subtle, but persistent, transcriptional signature indicative of independent origins. To explore this possibility, we separated the endoderm cells according to their lineage to identify differentially regulated genes that closely track with their developmental history. Strikingly, two X-linked genes, Trap1a and Rhox5, are consistently up-regulated in the extra-embryonic origin endoderm across lineage-traced embryos (K–S test, Bonferroni corrected p-value <0.05, **Fig. 5d, e**). Both genes are general markers for extra-embryonic tissue^25,26^, supporting a model where these cells diverge prior to specification of the embryo proper. Additionally, in an independently generated scRNA-seq data set of mouse embryogenesis, these markers exhibit uniform expression in extra-embryonic endoderm and extra-embryonic ectoderm, but highly heterogeneous expression in hindgut (**Fig. S8**)^10^. These observations corroborate our lineage-based evidence that a subpopulation of these seemingly embryonic endoderm cells derive from an extra-embryonic origin. We conclude that our lineage tracer can successfully pinpoint instances of convergent transcriptional regulation during embryogenesis.

### Towards a quantitative fate map

Simultaneous single cell recording of historical lineage and phenotype also provides the unique opportunity to infer the cellular potency and specification biases of ancestral cells reconstructed in our fate map. For example, we can set a lower boundary on the progenitor field size by counting the number of nodes in the tree that produce multiple tissue types (**Methods**).

Each node, which represents a unique lineage identity, can be thought of as a single cell that is reconstructed by our technology. In this manner, we investigated the founding number of progenitors during the earliest transitions in cellular potential. We defined totipotency as a node that gives rise to both embryonic and extra-embryonic ectodermal/placental cell types and tiered pluripotency into “early” and “late” according to the ability to form extra-embryonic endoderm, which is restricted independently from the epiblast within the inner cell mass (**Fig. 5f**). We find that the contributions of these founders to extant lineages are largely asymmetric, suggesting that even though a progenitor may be biased towards a specific fate, it retains the ability to contribute to other cell types. These results highlight the temporal disconnect between positional specification and true fate commitment and are expected in organisms that display indeterminate development. However, it is also possible that asynchronous expansion of a certain tissue type occurs after commitment, leading to substantial drift that could affect our inference of specification bias. Nonetheless, our data also provides lower bound estimates for the number of progenitors at these early developmental restriction points, suggesting a range of 1–6 totipotent cells, 4–17 early, and 15–52 late pluripotent progenitors from our current data (**Fig. 5g**). The variable number of multipotent cells at these stages reflects the encoded robustness that must exist to ensure successful assembly of a functioning organism, particularly given that only one pluripotent cell can be used to experimentally generate all somatic lineages in an embryo^27^.

## Discussion

In this study, we present a cell fate map underlying mammalian gastrulation using a custom technology for high information, multi-channel, and continuous molecular recording. Several key ideas have emerged, including the transformative nature of CRISPR-Cas9-directed mutation with a single cell RNA-seq read out^15–17^, how historical information from such a molecular recorder can complement RNA-seq profiles to characterize cell type, and an early framework for quantitatively understanding stochastic transitions during mammalian development.

The modularity of our molecular recorder allows for simple substitutions that will increase its breadth of applications. Here, we use three constitutively expressed guide RNAs of varying efficiencies to record continuously over time, but future experiments could employ environmentally-responsive promoters that sense stress, neuronal action potentials, or cell-to-cell contacts^28^, or combine these approaches for multifactorial recording. Similarly, Cas9-derived base editors^29^, including those that create diverse mutations^30^, can be substituted for the Cas9 nuclease, allowing for content-recording in cells that are particularly sensitive to DNA double strand breaks^31,32^.

Our cell fate map identifies an intriguing phenotypic convergence of independent cell lineages, showcasing the power of unbiased organism-wide lineage tracing to separate populations that appear similar in scRNA-seq alone. Specifically, we uncovered the surprising extra-embryonic origin of a subset of cells that resemble embryonic endoderm. While the initial specification of these two lineages are known to rely on largely redundant regulatory programs, they are temporally separated by several days, emerge from transcriptionally and epigenetically distinct progenitors, and form terminal cell types with highly divergent functions. The exact location or function of these extra-embryonic-originating cells is unclear, but the identification of highly predictive markers that segregate by origin provides a clear outline for further exploration. For example, previous studies have observed Trap1a expression within the embryonic endoderm^33^, suggesting that this population provides a novel developmental function despite its early spatial sequestration, and its precise location can be elucidated through spatial transcriptomics^34,35^. By showing the nearly complete convergence of cells of extra-embryonic origin to an embryonic endoderm state, our studies provide a functional foundation for classic lineage tracing experiments, which indicated a latent extra-embryonic contribution to the developing hindgut at the onset of gastrulation^36^. More generally, our approach can be used to investigate other convergent processes or to discriminate heterogeneous cell states that arise due to transient effects from persistent signatures that reflect a cell’s past, both of which will be critical for the complete assembly of a comprehensive cell atlas^37^. The scope of transdifferentiation within mammalian ontogenesis remains largely unexplored, but can be practically inventoried using our experimental system.

Ultimately, our cell fate mapping technology is designed to quantitatively address previously opaque questions in ontogenesis. Higher order issues of organismal regulation, such as the location, timing, and stringency of developmental bottlenecks, as well as the corresponding likelihoods of state transitions to different cellular phenotypes, can be modeled from the unbiased assembly of historical relationships. To that end, we used our platform to estimate the number of several multipotent progenitor fields and described the asymmetric biases of their developmental outcomes. Our hope is that characterization of these attributes will lead to new insights that connect large-scale developmental phenomena to the molecular regulation of cell fate decision-making.

## Methods

### Plasmid design and construction

Because the principles governing Cas9 efficiency and subsequent indel generation are not absolute, we screened fourteen protospacers for potential inclusion in our target site, including nine protospacers known to function with moderate efficiency and five additional protospacers hypothesized to function^1–10^. Each protospacer was checked against the human and mouse genomes using bowtie to limit off target effects. A gene block library of the fourteen protospacers (no additional bases between sequences) with an 8 base pair randomer was ordered from IDT representing target site version 0.0.

The target site (tS) v0.0 vector backbone was derived from a previously described Perturb-seq lentiviral vector (pBA439, Addgene, Cat#85967)^11^ with the following changes: the cassette for mU6-sgRNA-EF1a-PURO-BFP was removed and replaced with EF1a-tSv0.0-sfGFP using Gibson assembly with the target site in the coding sequence of sfGFP for use in the fluorescent reporter assay (PCT10, sequence available upon request).

A gene block library of five protospacers (ade2-whiteL-bam3-bri1-whiteB; no additional bases between sequences) with an 8 base pair randomer was ordered from IDT representing target site version 0.1. Protospacers in positions 1 (ade2), 3 (bam3), and 5 (whiteB), are used for cutting in subsequent experiments and are referred to as sites 1, 2, and 3.

Target site (tS) v0.1 was also cloned into pBA439 with the following changes: the cassette for mU6-sgRNA-EF1a-PURO-BFP was removed and replaced with EF1a-sfGFP-tSv0.1, followed by BGH pA on the original backbone (PCT12). Here the target site sits in the 3’ UTR of GFP. To improve the delivery of multiple targets into the same cell, we swapped the v0.1 target site cassette into a commercially available piggyBac transposon vector (Systems Biosciences, #PB533A-2) with the following changes: IRES-Neo was swapped for either GFP (PCT16) or mCherry (PCT29). The backbone was digested with restriction enzymes and target site v0.1 gene block was PCR-amplified to add Gibson arms. Following Gibson assembly, the plasmids were transformed into at least 100uL of Stbl2 competent cells (Thermo Fisher, Cat#10268019), and plated onto 1-2 large plates (Fisher, #NC9372402) with LB/Carbenicillin to generate high complexity target site libraries (PCT17, and PCT30, respectively).

The three-guide expression vector design and cloning protocol were adapted from ^11^ to utilize guides against the three sites in the target site. The guide for site 1 (ade2) is under the control of the mU6 promoter, site 2 (bam3) under the control of hU6 promoter, and site 3 (ade2) under the control of bU6-2 promoter. All guides are constitutively expressed in this system. Additionally, the triple-guide cassette was moved onto the piggyBac backbone described above.

Two further modifications of the plasmids described above were used in this study. First, in an attempt to decrease the cutting percentage variation between embryos, we cloned the triple-guide expression cassette without BFP into PCT29, and then cloned in the target site with intBCs to generate the resulting vectors (PCT41-43, for guide combinations (P,1,P), (1,1,1), and (2,1,2), respectively). In the second modification, we changed the truncated form of Ef1a in PCT29 to a promoter sequence comprised of the ubiquitous chromatic opening element (UCOE) and a full-length, intron-containing Ef1a and cloned in a triple-guide expression cassette for the guide combination (2,1,P), followed by cloning in of the target site to make PCT60. In these modifications, target site plasmid libraries (PCT41-43, PCT60) were transformed and expanded in 1-2L of liquid LB/Carb culture rather than on large plates.

### Cell culture, DNA transfections, and viral production

The production of lentiviral particles or transfection of plasmids as is as described in ^11^.

### K562 GFP reporter assay

To construct the target site GFP reporter cell line, a doxycycline(Dox)-inducible Cas9 K562 cell line was stably transduced with PCT10 (8% infected, <0.1 MOI), and GFP positive cells were sorted using fluorescence activated cell sorting on a BD FACSAria2. For each protospacer in the target site, 1-4 guides was designed to achieve a series of mutation efficiencies and cloned into single guide expression vectors^12^. On Day -4, the reporter cell line was plated into wells and stably transduced with a different guide against target site v0.0, GFP-targeting protospacer EGFP-NT2 (positive control), or Gal4-targeting protospacer (negative control) in each well. On Day -2, cells were selected for guide cassette integration using 3 ug/mL puromycin. On Day 0, 50ng/mL Dox was added to induce Cas9 expression, and maintained through the course of the experiment. GFP fluorescence was recorded on a LSR-II flow cytometer (BD Biosciences) on every 2^nd^ day starting at day 0, except day 13 was recorded in place of day 12. Data was analysed in Python using FlowCytometryTools (http://eyurtsev.github.io/FlowCytometryTools/). For guide virus produced in this experiment, labels were systematically shifted during production resulting in incorrect ordering of guide effect on GFP fluorescence, which was corrected for presentation in the manuscript. We confirmed the activity order of the guide series for three guides (ade2, bam3, and bri1) in sequencing experiments where new virus was prepared.

### K562 single cutting pooled assay

To construct the cell line used here, a Dox-inducible Cas9 K562 cell line was stably transduced with PCT12 (6% infected, <0.1 MOI), and GFP positive cells were sorted on a BD FACSAria2. On Day -5, the cell line was plated and stably transduced with a different guide against target site v0.1, or GFP-targeting protospacers in each well. On Day -2, cells were selected for guide cassette integration using 3 ug/mL puromycin. On Day 0, 50ng/mL Dox was added to induce Cas9 expression, and maintained through the course of the experiment. Wells were sampled every 3-6 days for 20 days with cell pellets frozen down. Genomic DNA was isolated from frozen cell pellets, and the target site was PCR-amplified to make sequencing libraries (refer to **Pooled embryo library preparation** below for library prep protocol), which were sequenced on the Illumina Miseq. Timepoint samples were pooled and reads with no indels were removed to calculate relative indel frequencies.

### K562 multiple target site integration cell line

To construct a cell line with multiple integrations, we nucleofected 200,000 Dox-inducible Cas9 K562 cells with 1500ng PCT17 and 200ng piggyBAC transposase using set program T-016 (Lonza #V4SC-2096; Systems Biosciences, #PB210PA-1).

### K562 triple guide cutting assay, and multi-clonal lineage tracing experiment

Multiple-integration cells described above were stably transduced with a triple guide expression vector (Perfect-Perfect-Perfect; fastest cutting) and recovered for 2 days. GFP (target site) and BFP (triple guide) double positive cells were sorted using fluorescence activated cell sorting on a BD FACSAria2. For the multi-clonal lineage tracing experiment, 10 cells were sorted into wells containing 200uL of pre-conditioned media on a 96 well plate (12 wells total). At day 18, wells were inspected under the microscope and the 3 wells with the largest populations were selected for single cell analysis on the 10x Chromium. Two of the lanes suffered wetting failures, and the remaining sample was taken through library preparation described below (refer to **Target site amplicon library preparation**). The library was sequenced on the Illumina Miseq and would benefit from additional sequencing.

For the pooled experiment, ~112,000 cells were sorted into a tube, spun down, resuspended in fresh media, split into two wells with 50ng/mL Dox added to one of the wells. Cells were collected 6 days post-sort, genomic DNA was isolated, and the target site was PCR-amplified to make sequencing libraries (refer to **Pooled embryo library preparation**), which were sequenced on the Illumina Miseq. The 10 intBCs with the most reads were used for analysis.

### Embryo generation

To enable *in vivo* lineage tracing, B6D2F1 strain female mice (age 6 to 8 weeks, Jackson Labs) were superovulated by sequential intraperitoneal injection of Pregnant Mare Serum Gonadotropin (5IU per mouse, Prospec Protein Specialists) and Human Chorionic Gonadotropin (5IU, Millipore) 46 hours apart. Twelve hours after delivery of the second hormone, MII stage oocytes were isolated and injected with *in vitro* transcribed piggyBAC transposase mRNA (100 ng/ul) prepared in an injection buffer (5 mM Tris buffer, 0.1 mM EDTA, pH = 7.4). Decapitated sperm isolated from an 8 week old *Gt(ROSA)26Sortm1.1(CAG=cas9*,EGFP)Fezh/J* strain mouse (Jackson labs, ref 13) was resuspended with the purified piggyBAC library in the same injection buffer at concentrations ranging from 0.5 to 1.4 ug/uL. Transposase-injected oocytes were then fertilized by piezo-actuated intracytoplasmic sperm injection (ICSI) as previously described ref 14. Injected embryos were cultured in 25 uL EmbryoMax® KSOM drops (Millipore) covered in mineral oil (Irvine Scientific) in batches of 25-50 embryos. After 84 or 96 hours, successfully cavitated blastocysts were screened for uniform fluorescence of the target sequence cassette and transferred into one uterine horn of 6-10 week old pseudopregnant CD-1 strain female mice (Charles River). Uterine transfer results in an ~24 hour lag, so the day of transfer was scored as E2.5 and embryos were dissected from euthanized animals 6 or 7 days later at ~E8.5 or E9.5, depending on the experiment. All techniques utilized standard micromanipulation equipment, including a Hamilton Thorne XY Infrared laser, Eppendorf Transferman NK2 and Patchman NP2 micromanipulators, and a Nikon Ti-U inverted microscope. Protocols are adapted from those described in ref 15.

### Pooled embryo library preparation

RNeasy Mini Kit (Qiagen, #74104) was used to isolate RNA from whole embryos or dissected tissue. Following first strand synthesis of 1ug RNA with oligo dT and AMV RT (Promega), the target site was amplified using a 2-stage PCR protocol. In the 1^st^ stage, <100ng of diluted cDNA template, 0.6 uM MC38 (CGTCGGCAGCGTCAGATGTGTATAAGAGACAGTGCAGGAGCGGATTGCTTCGAACC), and 0.6 uM MC39 (TCTCGTGGGCTCGGAGATGTGTATAAGAGACAGACAACCACTACCTGAGCACCCAGTC) was amplified using Kapa HiFi HotStart ReadyMix according to the following PCR protocol: (1) 98C for 3 min, (2) 98C for 30 s, 69C for 30 s, 72C for 15 s (16 cycles), (3) 72C for 5 min. Following 0.7X SPRI selection, the elute served as template for 2^nd^ stage PCR, with 0.6uM barcoded P5 (AATGATACGGCGACCACCGAGATCTACAC[ILLUMINA INDEX]*TCGTCGGCAGCGTC*AGATGTGTA), 0.6uM barcoded P7 (CAAGCAGAAGACGGCATACGAGAT[ILLUMINA INDEX]*GTCTCGTGGGCTCGG*AGATGTGTATAAG), amplified using Kapa HiFi HotStart ReadyMix according to the following PCR protocol: (1) 98C for 3 min, (2) 98C for 30 s, 60C for 30 s, 72C for 30 s (4-6 cycles), (3) 72C for 5 min. PCR products underwent 0.6X SPRI-selection and were eluted in 20-40uL of elution buffer to produce the final library. Libraries were sequenced on the Illumina HiSeq 2500 (Rapid Run) or Miseq, with the following run parameters: Read 1: 175 cycles, i7 index: 8 cycles, i5 index: 8 cycles, Read 2: 175 cycles.

### Single cell embryo dissociation

Embryos are washed through several drops of PBS after isolation to reduce debris and put into ~100 uL PBS droplets on a microscope slide and screened for uniform fluorescence of the target site cassette on an Olympus IX71 inverted microscope running Metamorph. Selected embryos were dissociated to single cell suspensions by adding 100 uL of TrypLE (Invitrogen, #12605010) and pipetting the embryo or embryo pieces every 5 minutes for ~30 minutes until complete dissociation was visually confirmed. Trypsin was deactivated by adding 100 uL PBS+BSA is added to the droplet and moving cells into a 1.5 mL eppendorf tube, followed by several rounds of additional collection with 100 to 200 uL of PBS+BSA to a final volume of 1 mL. The dissociated cells are filtered through a Flowmi filter tip (Bel-Art Products, #H13680-0040) into a new tube, and spun down for 5 minutes at 1200 rpm on a tabletop centrifuge. Following the spin, 900uL of PBS+BSA is removed and the remaining volume is resuspended with an additional 900uL of PBS+BSA. The suspension is spun for 5 minutes at 1,200 rpm, 800 uL of PBS+BSA is removed, the remaining volume is spun for 5 minutes at 1,200 rpm, and PBS+BSA is removed until only ~30 uL of volume remains. 2 uL of the final resuspended cells were used for counting using a hemocytometer. We load ~17,000 cells into the 10x machine (Chromium Single Cell 3’ Library & Gel Bead Kit v2) for a targeted recovery of 10,000 cells.

### scRNA-seq library preparation and sequencing

Single cell RNA-seq libraries were prepared according to the 10x user guide, except for the following modification. After cDNA amplification, the cDNA pool is split into two fractions. 15uL of EB buffer is added to one of the fractions of 20uL of the cDNA pool, and scRNA-seq library construction proceeds as directed in the 10x user guide. RNA-seq libraries were sequenced on the Illumina HiSeq 4000 system.

### Target site amplicon library preparation

The target site-specific amplification protocol was adapted from ^11^. 50-100 ng of template from the cDNA pool, 0.3 uM P5-truseq-long (AATGATACGGCGACCACCGAGATCTACACTCTTTCCCTACACGACGCTCTTCCGATCT), 0.6 uM MC63 (TCTCGTGGGCTCGGAGATGTGTATAAGAGACAGTGCAGGAGCGGATTGCTTCGAACC) was split across four parallel PCR reactions, and was amplified using Kapa HiFi HotStart ReadyMix according to the following PCR protocol: (1) 95C for 3 min, (2) 98C for 15 s, then 69C for 15 s (8-12 cycles). Reactions were re-pooled during 0.9X SPRI selection, and eluted into 60 uL. A second PCR with the elute as the template, 0.3 uM P5 (AATGATACGGCGACCACCGA), 0.6 uM barcoded P7 (CAAGCAGAAGACGGCATACGAGAT[ILLUMINA INDEX]GTCTCGTGGGCTCGGAGATGTGTATAAG) was split across four parallel PCR reactions, and amplified using Kapa HiFi HotStart ReadyMix according to the following PCR protocol: (1) 95C for 3 min, (2) 98C for 15 s, then 69C for 15 s (6 cycles). Reactions were re-pooled during 0.9X SPRI selection and then fragments of length 200-600bp were selected using the BluePippin. Target site libraries were sequenced on the Illumina HiSeq 2500 (Rapid Run), with the following run parameters: Read 1: 26 cycles, i7 index: 8 cycles, i5 index: 0 cycles, Read 2: 350 cycles.

### scRNA-seq library data processing

scRNA-seq data was processed and aligned using 10x Cell Ranger v2. The filtered gene-barcode matrices were then processed in Seurat (https://satijalab.org/seurat/) for data normalization (global scaling method “LogNormalize”), dimensionality reduction (PCA), and generation of t-sne plots, which use the first 16 principal components.

### scRNA-seq tissue assignment

An independent project conducting scRNA-seq profiling of gastrulation identified 42 distinct tissues in wild type mice. We were provided the mean expression profile for each tissue and the list of 712 marker genes used for assignment of cells to tissues. For each cell in lineage traced mouse embryos, we calculated the Euclidean distance between the cell’s expression profile and the mean expression profile for each tissue using the 712 marker gene set, and assigned the cell to the tissue identity with the minimum distance. Expression values were transformed to log space using log(normalized UMI count + 1) before calculating the Euclidean distance.

### Embryo gastrulation stage assignment

The wild type mouse gastrulation compendium consists of five time points, profiling every 0.5 days from E6.5 to E8.5 with at least 10 embryos collected for each time point. Tissue proportion is calculated as the number of cells assigned to the tissue divided by the total number of cells in the embryo. The median tissue proportion was calculated for each time point treating each tissue independently. For each lineage-traced embryo, the Euclidean distance between its tissue proportions and the median tissue proportion for each time point was calculated and the embryo was assigned to the time point with the minimum cumulative distance. All lineage-traced embryos were assigned to either E8.0 or E8.5 stages.

### Target site data processing

A custom software pipeline was built to align and call indels in the target site. The logic is as follows: (1) Assign cell barcode and UMIs to each read, (2) find the consensus sequence for each UMI, (3) align the consensus sequence to the target site reference sequence, (4) identify most likely integration barcodes (intBC) and create custom reference sequences, (5) repeat alignment against all reference sequences and select highest scoring alignment for each UMI, (6) call intBC and indels in the target site, (7) correct the intBC and allele using UMIs which appear in the same cell, (8) remove doublets. Details appear below:

Assigning cell barcode and UMIs to each read. Specific amplification libraries of the target site amplicon were processed using 10x Cell Ranger software to assign cell barcodes and UMIs to each read. The target site is designed to be orthogonal to the human and mouse genome, and does not align in Cell Ranger processing. Unaligned reads from the Cell Ranger output bam file are parsed into fastq format with the cell barcode and UMI identifiers appended to the read name.

Finding the consensus sequence for each UMI. To potentially increase the speed of consensus sequence finding, we attempt to trim reads to the same length for each UMI. The read is trimmed to remove sequence beyond the polyA tail using cutadapt software (http://cutadapt.readthedocs.io/en/stable/) with the following parameters: [–a AAAAAAAAAA – e 0.1 –o trimmedFile.fq –untrimmed-output=untrimmedFile.fq –m 20 –max-n=0.3 –trim-n]. Reads that do not contain polyA sequence appear in the untrimmed file and are subjected to a second round of read trimming using a sequence which appears in the 3’ end of the target site assuming the sequence has not been deleted from DNA repair, with cutadapt run using the following parameters: [-a GCTTCGTACGCGAAACTAGCGT -e 0.1 -o trimmedFile2.fq -- untrimmed-output=untrimmedFile2.fq -m 20 --max-n=0.3 --trim-n --no-indels]. The adapter sequence used in the last round of trimming is then concatenated back on to the trimmed sequence to improve target site alignment in the next step. If >=60% of trimmed sequences for a given UMI are the same sequence, then the sequence is taken as the consensus sequence. Otherwise, a multiple sequence alignment is performed using BioPython and the consensus sequence is extracted from the alignment. Ambiguous bases are reported if there is <50% agreement for any position in the alignment.

Aligning to the target site reference. We use the emboss implementation of the smith-waterman algorithm to align sequences to the target site reference sequence with the following parameters, which were determined empirically: [emboss water –asequence targetSiteRef.fa – sformat1 fasta –bsequence consensusUMI.fa –sformat2 fasta –gapopen 15.0 –gapextend 0.05 – outfile sam –aformat sam]. In this first alignment, the ambiguous sequence NNNNNNNN is used to represent the intBC. A minor bug had to be corrected in the emboss implementation to successfully output sam format.

Identifying the most likely intBC. A perl script is used to parse the intBC from the alignment. The intBCs with the highest number of UMIs are substituted into the target site reference sequence to make custom reference sequences. This step was included because upon manual inspection, there were obvious misalignments due to the ambiguous intBC sequence, which were corrected upon substitution of a real sequence.

Selecting the highest scoring alignment for each UMI. Repeat smith waterman alignment against all custom reference sequences and select alignment with the highest score for each UMI.

Calling indels and intBCs. A perl script is used to parse the intBC and indels from the alignment using the CIGAR string. The boundaries for each site is defined and indels overlapping site boundaries are called as an indel in that site. Sequence of the indel is not considered.

Correcting indels using multiple reads with the same UMI from the same cell. UMIs are filtered for alignment score and only cells that are in the matched scRNA-seq data set are kept. An intBC is corrected to an intBC with a higher UMI count in the cell if the intBCs are within an edit distance of 2 and the alleles are the same. An allele is the combination of indels in sites 1, 2, and 3. An allele is corrected to an allele with a higher UMI count in the cell if the intBC is the same and the allele is within a 1-indel difference. Only UMIs past a minimum read count threshold are kept.

Eliminating doublets. Cells that report two alleles for the same intBC are removed if the dominant allele is <80% of the total UMI count for the intBC. This removes 4.1-18.3% of cells in our embryos.

### Tree reconstruction strategy

We simulate the evolutionary process leading from a collection of uncut target sites to the final data set. The set of mutations (including “no mutation”) across all target sites in a cell is referred to as an allele. In the final tree, each branch represents a mutation, and each node represents the allele of a cell, which may be a reconstructed ancestral allele, i.e. it is not present in the data set.

Input: table of unique alleles

- each allele may represent multiple cells
- we cannot distinguish between identical indels in the same position that may result from independent mutation events (convergent indels) if they appear with an identical set of co-segregating indels

Algorithm:

- Create root node in tree representing an allele with 0 mutations (c_allele)
- remove alleles in the table that match c_allele
- While alleles remain in table:
  - choose indel from table that can be added to current allele
  - can only add indels in positions that have no mutation
  - create new node by adding indel into c_allele (c_allele2)
  - draw directed edge labeled with indel between nodes from c_allele to c_allele2
  - remove alleles in table that match c_allele2
  - includes alleles that match c_allele2 with missing values for positions that have no mutations
  - if indels in table can be added to c_allele2, then c_allele = c_allele2; else, c_allele does not change
  - when indels cannot be added to c_allele, traverse up edges to ancestral nodes until an allele to which an indel can be added is found

We presented two methods that are used to choose indels. The first method, “Random”, selects a position where an indel can be added, and then selects an indel from the data set for that position; both selections occur in an unbiased manner. The second method, “Frequency Normalized Weighted” (FNW), identifies all of the indels that can be added to the current allele and scores them according to the fraction of alleles they are found in divided by the expected independent frequency of the indel (see **Figure 2d**). These scores are used as weights to bias selection of the indel. The reasoning behind FNW is that indels that are found in many cells (or alleles) are more likely to have occurred early, but this has to be balanced against their expected likelihood of occurring. FNW biases the search towards more likely trees. To further increase the search for good trees, we first remove all indels that are unique to a single allele since we can assume that these indels occur at the leaves of the tree. The indels are added as branches leading to leaves in the final tree before tree likelihood is calculated.

The log likelihood of the tree is calculated as the sum of the likelihoods of all the indels that appear in the tree. The likelihood of each mutation is estimated from the embryo data set (**Figure 2d**).

It is worth noting that the number of trees that are possible grossly exceeds 30,000; however, the search is biased towards finding good trees and performs markedly better than those that are randomly generated. Using high scoring trees to direct the search towards better ones, such as by grafting high scoring branches, could further improve our algorithm’s ability to identify high scoring trees.

Trees are visualized using the Python ete library (http://etetoolkit.org/).

### Estimating ancestral tissue relationships

Each node, including leaves, that includes more than one tissue type is considered a “progenitor.” Progenitors found at the leaves are not reconstructed or inferred but result from the lack of new indels that distinguish between tissues (ie the lineage tracer does not produce new indels past the progenitor stage). For reconstructed progenitors, only nodes with >0.8 median branch support across similarly high scoring trees are included in this analysis (See **Fig. S7**).

The shared progenitor score is calculated between two tissues as the number of shared progenitors scaled by the number of tissues each progenitor contributes to, and is calculated using the following algorithm:

For each progenitor,

> tList = list of tissues progenitor contributes to pScore = 1/len(tList)
>
> for each pair of tissues in tList: progenitorScoreForPairOfTissues += pScore
>
> Example for a single progenitor: tList = [Endo, Meso, XMeso] pScore = 1/3
>
> ProgenitorScoreEndoMeso += 1/3 ProgenitorScoreEndoXMeso += 1/3 ProgenitorScoreMesoXMeso += 1/3

Example for a single progenitor:

> tList = [Endo, Meso, XMeso]
>
> pScore = 1/3
>
> ProgenitorScoreEndoMeso += 1/3
>
> ProgenitorScoreEndoXMeso += 1/3
>
> ProgenitorScoreMesoXMeso += 1/3

The resulting matrix is a shared progenitor score matrix. To transform the similarity matrix to a distance matrix, we use 1-(matrix/maxScoreInMatrix). The distance matrix is then hierarchically clustered with centroid joining.

### Endoderm lineage assignment and differentially regulated gene identification

Endoderm cells can have one of three origins based upon our tree: extra-embryonic, embryonic, or ambiguous. Cells are considered extra-embryonic if there is a progenitor in its lineage whose descendants include ≥ 40% extra-embryonic cells. Cells have ambiguous origin if they descend directly from the root node. Otherwise, cells are considered to be from embryonic origin. We identified endoderm cells of extra-embryonic origin in all embryos but embryo 7.

We use the Kolmogorov-Smirnov test (Python scipy.stats.ks_2samp) to identify differentially regulated genes between embryonic and extra-embryonic origin endoderm cells. Only highly variable genes in the embryo are considered for testing, and genes are significant if they have a Bonferroni corrected p-value under 0.05.

### Multipotent field size estimation and asymmetry

Progenitors are considered pluripotent if their descendants include at least one mesoderm (Meso or XMeso or Blood) cell, one ectoderm (Ecto) cell, and one endoderm (Endo) cell. A pluripotent progenitor are considered early pluripotent if it also has at least one extra-embryonic endoderm descendant, and further considered totipotent if it has at least one extra-embryonic ectoderm descendant. To estimate the lower bound for the number of multipotent cells, we start at the bottom level of the tree and count the number of multipotent cells at that level. If multipotent cells exist, then the number of multipotent cells is propagated to its ancestor in the above level, otherwise we count 0 for that level. We add one progenitor for every level that includes a multipotent cell and other cells to represent the progenitor that lead those non-multipotent cells at that level. The number of multipotent cells is then the number of cells propagated to the root of the tree. Progenitor asymmetry is simply the proportion of descendants from each of the tissues for that node.

## Acknowledgments

The authors would like to thank members of the Weissman and Meissner labs for their helpful discussions and insight towards the production of this work, as well as Eric Chow and Derek Bogdanoff of the UCSF CAT for their help with sequencing. This work was funded by National Institutes of Health Grants R01 DA036858 and 1RM1 HG009490-01 (J.S.W.), P50 HG006193 and R01 HD078679 (A.M.), F32 GM116331 (M.J.), and F32 GM125247 (J.J.Q), and Chan-Zuckerberg Initiative 2018-184034. J.S.W. is a Howard Hughes Medical Institute Investigator. M.M.C. is a Gordon and Betty Moore fellow of the Life Sciences Research Foundation. T.M.N. is a fellow of the Damon Runyon Cancer Research Foundation. S.G., H.K., and A.M. are supported by the Max Planck Society.

## Author Contributions

M.M.C., Z.D.S., A.M. and J.S.W. were responsible for the conception, design, and interpretation of the experiments and wrote the manuscript. M.M.C. and Z.D.S. conducted experiments and developed the analysis, with input from Z.D.S. S.G. and H.K. provided annotations for RNA-seq data. B.A., T.M.N, and M.J. provided vectors, experimental protocols, and advice from perturb-seq, from which the recorder technology is developed. J.Q. and D.Y. were engaged in discussion of the technology.

## Author Information

Data will be deposited into GEO prior to publication. The authors declare no competing interests.

